# Dissecting the Role of the Human Microbiome in COVID-19 via Metagenome-assembled Genomes

**DOI:** 10.1101/2022.03.09.483704

**Authors:** Shanlin Ke, Scott T. Weiss, Yang-Yu Liu

**Author notes:** Correspondence and requests for materials should be addressed to Y.-Y.L.

## Abstract

Coronavirus disease 2019 (COVID-19), primarily a respiratory disease caused by infection with Severe Acute Respiratory Syndrome Coronavirus 2 (SARS-CoV-2), is often accompanied by gastrointestinal symptoms. However, little is known about the relation between the human microbiome and COVID-19, largely due to the fact that previous studies fail to provide high taxonomic resolution to identify microbes that likely interact with SARS-CoV-2 infection. Here we used whole-metagenome shotgun sequencing data together with assembly and binning strategies to reconstruct metagenome-assembled genomes (MAGs) from a total of 514 nasopharyngeal and fecal samples of patients with COVID-19 and controls. We reconstructed a total of 11,584 medium-and high-quality microbial MAGs and obtained 5,403 non-redundant MAGs (nrMAGs) with strain-level resolution. We found that, thanks to the high taxonomic resolution of nrMAGs, the gut microbiome signatures can accurately distinguish COVID-19 cases from healthy controls and predict the progression of COVID-19. Moreover, we identified a set of nrMAGs with a putative causal role in the clinical manifestations of COVID-19 and revealed their functional pathways that potentially interact with SARS-CoV-2 infection. The presented results highlight the importance of incorporating the human gut microbiome in our understanding of SARS-CoV-2 infection and disease progression. The genomic content of nrMAGs presented here has the potential to inform microbiome-based therapeutic developments for COVID-19 progression and post-COVID conditions.

## Introduction

The ongoing pandemic of coronavirus disease 2019 (COVID-19), a respiratory disease caused by severe acute respiratory syndrome coronavirus 2 (SARS-CoV-2), has infected billions of people world-wide. A broad range of clinical manifestations of COVID-19 has been reported, including asymptomatic or mild disease with cough and fever to severe pneumonia with multiple organ failure and acute respiratory distress syndrome (ARDS) leading to death^1^. Existing studies found that a large proportion of COVID-19 patients had at least one gastrointestinal (GI) symptom^2-5^, such as diarrhea, vomiting, or belly pain. Moreover, it has been reported that, among 73 SARS-CoV-2-infected hospitalized patients in China, 53.4% of patients tested positive for SARS-CoV-2 in their stool samples ranging from day 1 to 12 post infection^6^. Importantly, in more than 20% of infected patients, their fecal samples remained positive for the virus even after the respiratory and/or sputum samples exhibited no detectable virus^6^. In some cases, the viral load in feces is even higher than that in pharyngeal swabs^3^. All these results suggest that the GI tract might be an important extra-pulmonary site for SARS-CoV-2 infection. Currently, the role of angiotensin-converting enzyme 2 (ACE2) in the invasion of host cells by SARS-CoV-2 via its spike protein is well-established^7^, and ACE2 is also highly expressed in the small intestine and colon^4,8^. Therefore, the prolonged presence of large amounts of fecal SARS-CoV-2 RNA virus is unlikely to be explained by the swallowing of virus particles replicated in the throat but rather suggests enteric infection with SARS-CoV-2.

The human GI tract is the largest immune organ in the body and plays a critical role in the immune response to pathogenic infection or commensal intrusion^9^. Trillions of microbes live inside the GI tract. Those microbes and their genes, collectively known as the human gut microbiome, modulate host immunity^10^. To date, several studies, based on 16S rRNA gene sequencing, have demonstrated that the human upper respiratory and gut microbiome are broadly altered in patients with COVID-19^11-15^. Although 16S rRNA gene sequencing provides valuable insights into the general characteristics of the human microbiota, it does not offer the taxonomic resolution needed to capture sufficient sequence variation to discriminate between closely related taxa^16^. Other studies, based on whole-metagenome shotgun (WMS) sequencing, explored the links between the human microbiome and SARS-CoV-2 infection by mapping short metagenomic reads to reference genome databases^17-21^. Despite the fact that the analysis of WMS sequencing data provides more information than 16S rRNA gene sequencing data analysis, existing studies based on reference genome databases are subject to the limitations and biases of those databases and unable to characterize microbes that do not have closely related culture representatives. In fact, an estimated 40–50% of human gut species lack a reference genome^22,23^. This may result in a strong null bias in characterizing the gut microbial community.

An alternative strategy to analyze WMS sequencing data is to reconstruct metagenome-assembled genomes (MAGs) through *de novo* assembly and binning^24^. One key advantage of this strategy is that it allows recovery of genomes for microorganisms that have yet to be isolated and cultured and hence are absent from the current reference genome databases. This strategy has been adopted in several studies to provide genomic insights into microbial populations that are critical to human health and disease^25,26^. In this study, we applied state-of-the-art metagenome assembly and binning strategies to reconstruct microbial population genomes directly from microbiome samples of COVID-19 patients and controls (**Fig. 1**). Our major goals were to construct a COVID-19 related metagenomic genome catalog to identify novel taxa and strain-level differences that are likely related to the clinical manifestations of SARS-COV-2 infection. Our results demonstrate the association of the human microbiome and SARS-COV-2 infection at an unprecedentedly high level of taxonomic resolution. More importantly, our study provides a unique resource to directly investigate the genomic content of COVID-19 relevant microbial strains and sheds light on more targeted follow-up studies.

**Fig. 1.**
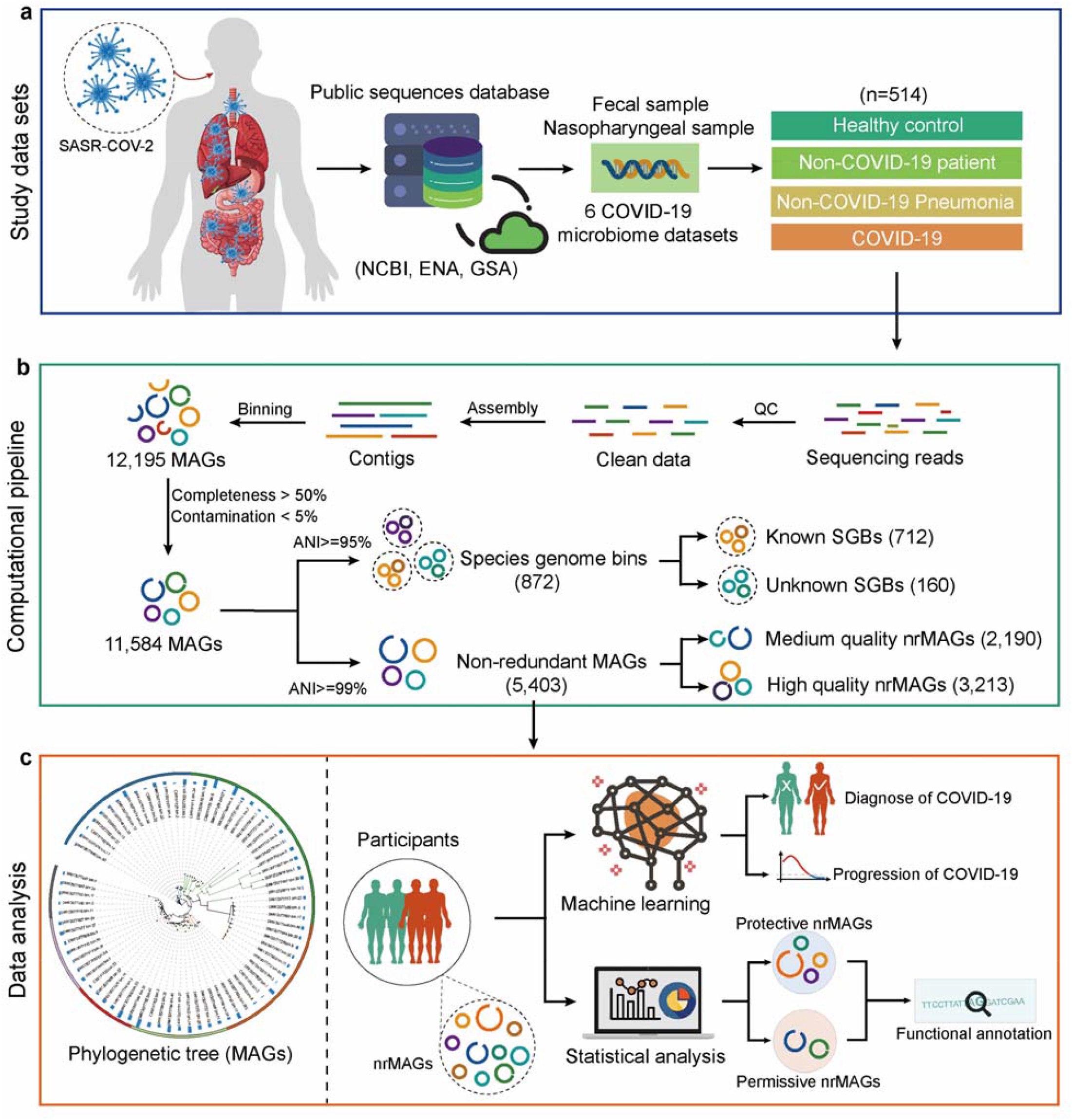
Conceptual framework of study. **a**, To understand the relation between human microbiome and COVID-19 via metagenome-assembled genomes (MAGs), we collected a total of 514 shotgun metagenomic sequencing data from 6 public data sets. These microbiome sample including fecal and nasopharyngeal samples from COVID-19 and non-COVID-19 controls. **b**, A total of 11,584 MAGs (≥50% completeness and ≤5% contamination) were constructed from all metagenomic sequencing data. The reconstructed MAGs were first clustered to 872 species-level genome bins (SGBs) at 95% of the ANI (average nucleotide identity). SGBs containing at least one reference genome (or metagenome-assembled genome) in the Genome Taxonomy Database (GTDB) were considered as *known* SGBs. Otherwise, they were considered as *unknown* SGBs. The reconstructed MAGs were then dereplicated to 5403 non-redundant MAGs (nrMAGs, strain level) based on 99% of ANI. The 5,403 nrMAGs were divided into medium-quality MAGs (50% ≤ completeness < 90% and ≤5% contamination) and high-quality MAGs (≥90% completeness and ≤5% contamination). **c**, The phylogenetic tree of nrMAGs was constructed using PhyloPhlAn. We employed Random Forest machine learning models together with nrMAGs to diagnose COVID-19 and predict the progression of COVID-19 (date of negative RT-qPCR results). The permissive and protective nrMAGs of COVID-19 were identified by GMPT pipeline. And the genomes of permissive and protective nrMAGs were functionally annotated using Prokka and MicrobeAnnotator.

## Results

### COVID-19 related metagenomic data sets

To examine the relation between the human microbiome and COVID-19 via MAGs, we first gathered WMS sequencing data from the COVID-19 related human microbiome studies (publicly available as of August 2021). We collected the raw WMS sequencing data of 514 microbiome samples (n=359 individuals) from six publicly available datasets (**Fig.1a** and **Table 1**) with different technical settings (e.g., sequencing platform and sequencing depth), including nasopharyngeal (n=96) and fecal microbiome (n=418) samples. Among these data sets and samples, we have 404 (78.60%) and 110 (21.40%) samples from COVID-19 and Non-COVID-19 controls (**Fig.2a**), respectively.

**Fig. 2.**
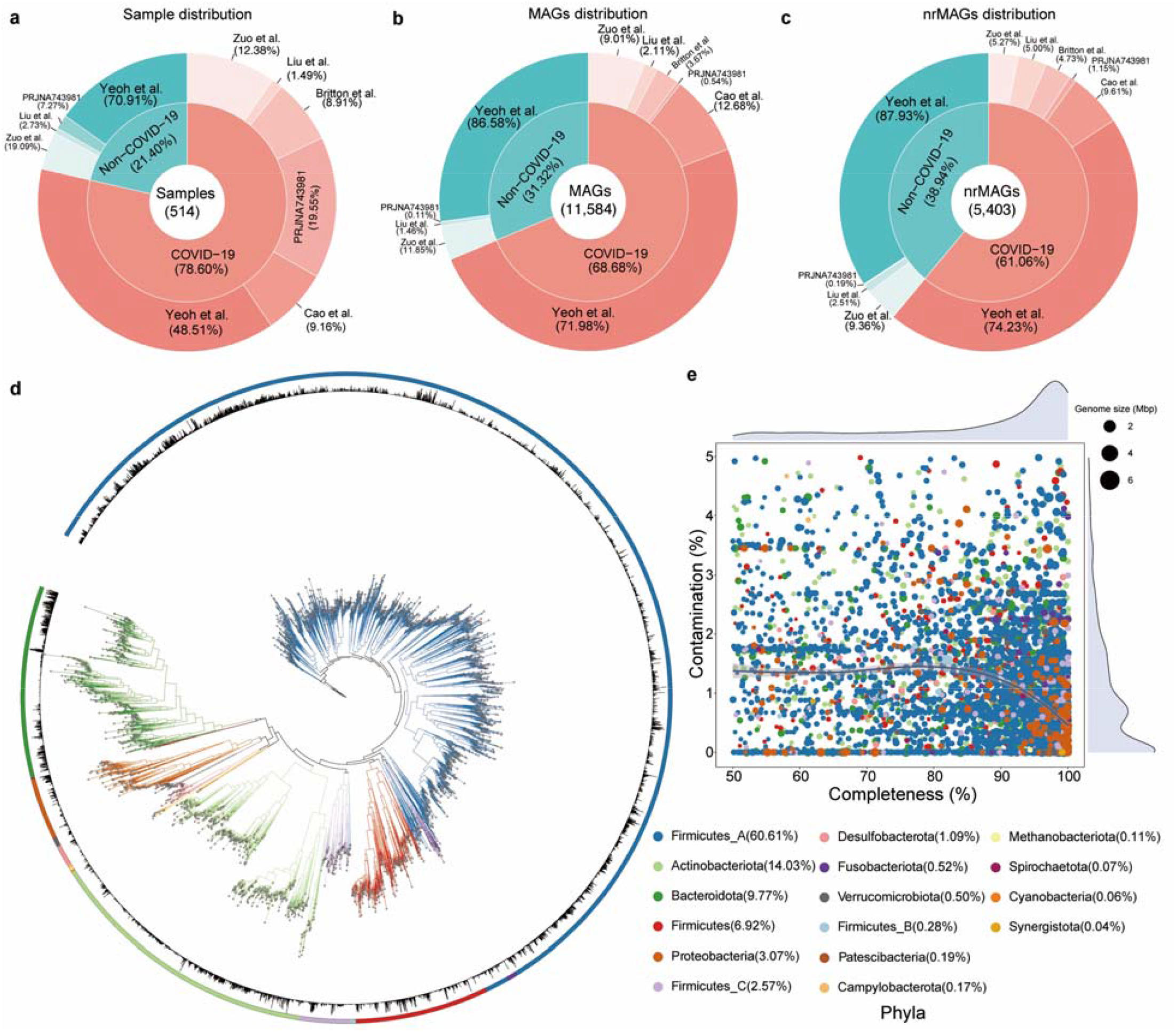
Reconstruction of MAGs from 514 COVID-19 related shotgun metagenomics sequencing data. **a**, Sample distribution among different dataset, source type, and disease status. **b**, Number of MAGs recovered from different dataset, source type, and disease status. **c**, Number of nrMAGs recovered from different dataset, source type, and disease status. **d**, Phylogenetic tree of nrMAGs constructed using PhyloPhlAn. The color of outer cycle and clades represents phylum and bar plot within cycle represents the average relative abundance across all microbiome samples. **e**, The distribution of completeness and contamination on nrMAGs and the color of point represents phylum. And the size of point represents the genome size of nrMAGs.

**Table 1.** Human metagenome data sets analyzed in this study.

| Dataset (year) | Zuo et al.<sup>17</sup> | Britton et al.<sup>53</sup> | Cao et al.<sup>54</sup> | Yeoh et al.<sup>18</sup> | Liu et al.<sup>35</sup> | PRJNA743981<sup>36</sup> | Total |
| --- | --- | --- | --- | --- | --- | --- | --- |
| COVID-19 positive samples | 50 | 36 | 37 | 196 | 6 | 79 | 404 |
| COVID-19 negative samples | 21 | 0 | 0 | 78 | 3 | 8 | 110 |
| Total subjects | 36 | 36 | 13 | 178 | 9 | 87 | 359 |
| Total samples | 71 | 36 | 37 | 274 | 9 | 87 | 514 |
| Longitudinal | Yes | No | Yes | Yes | No | No | - |
| Source | Fecal | Fecal | Fecal | Fecal | Nasopharyngeal | Nasopharyngeal | - |
| Geography | CHINA | USA | CHINA | CHINA | CHINA | - | - |
| Year | 2020 | 2021 | 2021 | 2021 | 2021 | 2021 | - |
| Sequencing platform | Illumina<br>Novaseq<br>6000 | Illumina<br>Hiseq | HiSeq<br>XTen<br>(PE250) or<br>NovaSeq<br>(PE150) | Illumina<br>Novaseq<br>6000 | Illumina<br>Novaseq 6000 | Illumina<br>Novaseq 6000 | - |
| Total sequences (mean) | 17,245,724.1<br>8 | 4,636,385.22 | 19,666,057.<br>43 | 23,717,063.9<br>9 | 40,593,115.44 | 28,683,197.33 | - |
| Accession number | PRJNA6242<br>23 | PRJNA66088<br>3 | PRJCA003<br>532 | PRJNA6502<br>44 | PRJNA656660 | PRJNA743981 | - |

#COVID-19 negative samples are healthy controls or Non-COVID-19 patients who tested negative for SARS-CoV-2 infection.

### A high-quality microbial genome catalog of COVID-19

After quality control, we performed metagenomic assembly and binning on those microbiome samples and recovered 12,195 MAGs in total (**Fig.1b**). To standardize the genome quality across all datasets, we used thresholds of ≥50% genome completeness and ≤5% contamination^27,28^, resulting in 11,584 MAGs [mean completeness□=□87.55%; mean contamination□=□0.99%; mean genome size =2.6 megabases (Mb); mean N50 =61.8 kilobases (kb), **Fig.S1**,**S2**]. Here N50 is the sequence length of the shortest contig at 50% of the total genome length. To obtain the view of the microbial community at the species level, we first organized 11,584 MAGs into species-level genome bins (SGBs) at an ANI (average nucleotide identity) threshold of 95%, resulting in a total of 872 SGBs, of which 160 (18.35%) SGBs represented species without any available genomes from the Genome Taxonomy Database (GTDB)^29^ and were defined as unknown SGBs (uSGB, **Fig.S3**). To evaluate the highest quality representative genomes, we dereplicated the 11,584 MAGs at an ANI threshold of 99%, resulting in a final set of 5,403 non-redundant MAGs (nrMAGs) with strain-level resolution [mean completeness□=□86.87%; mean contamination□=□0.99%; mean genome size =2.4 megabases (Mb); mean N50 =63.2 kilobases (kb), **Fig.2d-e and Fig.S2**]. We found that each Non-COVID-19 microbiome sample contributed relatively higher rates of total MAGs and nrMAGs than COVID-19 microbiome samples as 21.40% Non-COVID-19 microbiome samples contributed to 31.32% of total MAGs and 38.94% of nrMAGs (**Fig.2a-c** and **Fig.S4**).

Among those 5,403 strain-level nrMAGs, 2,190 (40.53%) nrMAGs satisfied the medium-quality criteria (50% ≤ completeness < 90% and ≤5% contamination), and 3,213 (59.47%) nrMAGs showed high-quality (≥90% completeness and ≤5% contamination) (**Fig.1**)^27,28^. Using the Genome Taxonomy Database^29^, 5,397 (99.89%) and 6 (0.11%) nrMAGs were assigned to bacterial and archaeal domains, respectively (**Fig.S5**). As shown in **Fig.2d**, the nrMAGs came predominantly from five phyla: the Firmicutes A (60.61%), Actinobacteriota (14.03%), Bacteroidota (9.77%), Firmicutes (6.92%), and Proteobacteria (3.07%). These dominant phyla identified from our study are quite consistent with that of prior work of the Human Gastrointestinal Microbiota Genome Collection (human gastrointestinal bacteria culture collection combined with publicly available, high-quality human gastrointestinal-associated bacterial genomes)^30^. Moreover, the archaeal nrMAGs were all taxonomically assigned to the archaeal phylum Methanobacteriota.

### Alterations of the human microbiome in COVID-19 patients

Previous studies demonstrated that SARS-CoV-2 infection is associated with the alpha diversity of the human gut^12,31,32^ and oral^33,34^ microbiome at the genus- or species-level. We first investigated if SARS-CoV-2 infection is associated with alpha diversity of the human microbiome at the nrMAG-level. The alpha diversity measures (i.e., Richness and Shannon index) from COVID-19 patients and Non-COVID-19 controls were compared (**Fig.3a** and **Fig.S6**). In accordance with previous studies, we found that the Richness and Shannon index of the gut microbiome in COVID-19 patients were significantly lower than that in Non-COVID-19 controls in two datasets (Zuo et al.^17^ and Yeoh et al.^18^, **Fig.3a** and **Fig. S6a**,**d**). Interestingly, no significant results were found between patients with COVID-19 Pneumonia and Non-COVID-19 Pneumonia controls (**Fig.3a** and **Fig. S6a**). Consistent with a previous study^35^, we found no significant differences between COVID-19 patients and Non-COVID-19 controls in the nasopharyngeal microbiome samples (**Fig.3a** and **Fig.S6e**,**f**). This may be due to the small sample size and the fact that Non-COVID-19 controls are not health controls in one of the nasopharyngeal microbiome datasets (Liu et al.^35^). Moreover, in the other nasopharyngeal microbiome dataset (PRJNA743981^36^), we only identified a small number of nrMAGs, as a large portion of sequencing reads from this dataset were contamination from the human genome.

**Fig. 3.**
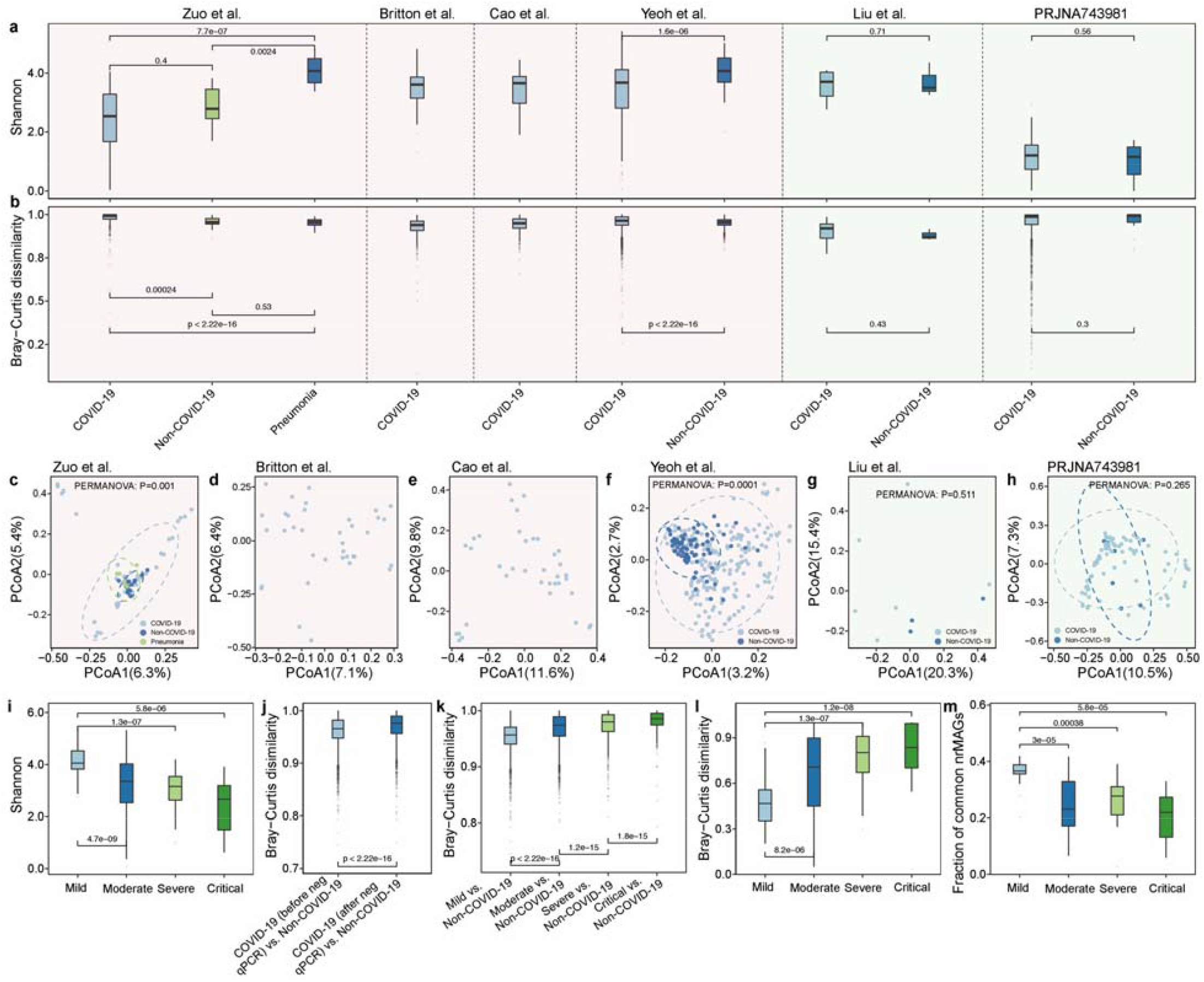
COVID-19 related alterations of the human microbiome. Shannon diversity (**a**) and within group Bray-Curtis dissimilarity (**b**) of the human microbiome at the nrMAG-level from each study. Principal Coordinates Analysis (PCoA) plot based on Bray–Curtis dissimilarity of microbial compositions from the study of Zuo et al. (**c**), Britton et al. (**d**), Cao et al. (**e**), Yeoh et al. (**f**), Liu et al. (**g**), and PRJNA743981 (**h)**. All PERMANOVA tests were performed with 9999 permutations based on Bray–Curtis dissimilarity. The background color of each panel (**a-h**) represents the source of the microbiome samples from the human gut (light red) or nasopharynx (light green). **i**, Shannon diversity at different disease severity groups. **j**, Boxplot of the gut microbiome Bray–Curtis dissimilarity between healthy controls and patients with COVID-19 before or after their nasopharyngeal aspirates or swabs tested negative for SARS-CoV-2 via RT-qPCR. **k**, Boxplot of the gut microbiome Bray–Curtis dissimilarity between healthy controls and patients with COVID-19 from different disease severity groups. **l**, Boxplot of Bray–Curtis dissimilarity of individual microbiome temporal changes over time from different disease severity groups. **m**, The fraction of common nrMAGs on patients with COVID-19 over time. *P* values were calculated by two-sided Wilcoxon–Mann–Whitney test.

In line with previous studies^11,37^, PCoA (principal coordinates analysis) combined with PERMANOVA (permutational multivariate analysis of variance) revealed that two datasets (from Zuo et al.^17^ and Yeoh et al.^18^, **Fig.3c-h**) had a significant difference in microbial community structure between the patients with COVID-19 and Non-COVID-19 controls at the nrMAG-level. Moreover, the patients with COVID-19 in these two datasets showed significant higher within-group variation than that in the Non-COVID-19 controls (**Fig.3b**).

In the data set from Yeoh et al.^18^, the gut microbiome samples from those patients with COVID-19 were collected before and after their nasopharyngeal aspirates or swabs tested negative for SARS-CoV-2 via RT-qPCR. Furthermore, patients with COVID-19 were classified into four severity groups (i.e., mild, moderate, severe, and critical) based on symptoms as reported in the previous study^38^. We found that patients with COVID-19 who had milder disease severity showed significant higher Shannon diversity in their gut microbiome (**Fig.3i**). Interestingly, the composition of nrMAGs in patients with COVID-19 after recovery (negative for SARS-CoV-2 via RT-qPCR) were significantly different from Non-COVID-19 controls than from patients with COVID-19 before recovery (positive for SARS-CoV-2 via RT-qPCR, **Fig.3j**. In line with a previous study at the metabolic capacity level^39^, these results indicate that the gut microbiome of patients with COVID-19 did not return to a relatively healthy status right after their recovery from SARS-CoV-2 infection. We then observed that disease severity of COVID-19 was significantly positively associated with the gut microbiome dissimilarity between COVID-19 patients and Non-COVID-19 controls (**Fig.3k**). To understand the relation between disease severity and short-term variation in the gut microbiome of patients with COVID-19, we traced the changes in the microbiome within each individual associated with disease severity. Interestingly, patients with COVID-19 who had milder disease severity showed lower temporal variation in the gut microbiome (quantified by the Bray-Curtis dissimilarity of longitudinal microbiome samples, **Fig.3l**). The lower temporal variation of gut microbiome samples in milder disease severity groups is partly due to the higher fraction of common nrMAGs across the longitudinal microbiome samples in those groups (**Fig.3m**).

### nrMAGs accurately classifies COVID-19 patients and Non-COVID-19 controls

Previous studies have demonstrated the diagnostic potential of the microbiome-based classification for SARS-CoV-2 infection using genus- or species-level taxonomic profiles^12,21,40^. To test whether the gut microbial composition at the nrMAG-level can distinguish COVID-19 patients from Non-COVID-19 controls, we built random forest classifiers on two datasets (Zuo et al.^17^: 50 patients with COVID-19 and 15 Non-COVID-19 controls; and Yeoh et al.^18^: 196 patients with COVID-19 and 78 Non-COVID-19 controls), separately. Importantly, this analysis was performed with 5-fold cross-validation and the data were randomly split into training and test sets 50 times. Since we had unbalanced classes, we applied two metrics to quantify the classification performance: AUROC (Area Under the Receiver Operating Characteristic curve) and AUPRC (Area Under the Precision-Recall Curve). Consistent with the PCoA analysis (**Fig.3c**), using the data from Zuo et al.^17^, we found that nrMAGs can accurately detect COVID-19 with the mean AUROC and AUPRC values of 0.981 and 0.971, respectively (**Fig.S7a**). The top COVID-19 related features included multiple nrMAGs from *Blautia A sp003480185, Blautia A wexlerae, Agathobacter faecis, Eisenbergiella sp900066775, Faecalibacterium prausnitzii G*, and *Lachnospira rogosae* (**Fig.S7b**). Consistent with the first dataset (Zuo et al.^17^) and the PCoA analysis (**Fig.3f**), the random forest classifier on the larger cohort (Yeoh et al.^18^) also showed high classification performance (AUROC∼0.920; AUPRC∼0.884; **Fig.S7c**). The key discriminatory nrMAGs of COVID-19 in this cohort belonged to *Adlercreutzia equolifaciens, Blautia A sp003471165, Eisenbergiella sp900066775, Eubacterium I, Gemmiger sp900539695, and Romboutsia timonensis* (**Fig.S7d**). Moreover, three specific nrMAGs were identified (from *Mediterraneibacter A butyricigenes* and *Eisenbergiella sp900066775)* as common features between the two data sets.

### nrMAGs accurately predict the progression of COVID-19

We next investigated the association between nrMAGs and the progression of COVID-19. To explore this association, we employed a random forest regression model to predict the date of negative RT-qPCR result using the data from Yeoh et al.^18^ (with 196 microbiome samples from 100 COVID-19 patients, **Fig.S8**). The regression tasks were performed with 5-fold cross-validation and we then randomly split the data 50 times. Remarkably, this approach demonstrated that the dates of negative RT-qPCR result were well predicted by nrMAGs (Pearson correlation 0.425, *P* value < 1e−30, **Fig.4a**). Among the top-30 (based on the percentage increase in mean squared error) most important nrMAGs (**Fig.4b**), we identified multiple species such as *Citrobacter freundii, Enterocloster sp900543885, Citrobacter portucalensis, Parabacteroides distasonis* and *Veillonella parvula*. Notably, we also found some nrMAGs from well-known opportunistic pathogens including MAG02074 (*Klebsiella quasivariicola*^41^), MAG03769 (*Klebsiella pneumoniae*^42^), and MAG02080 (*Escherichia coli D*^43^).

**Fig. 4.**
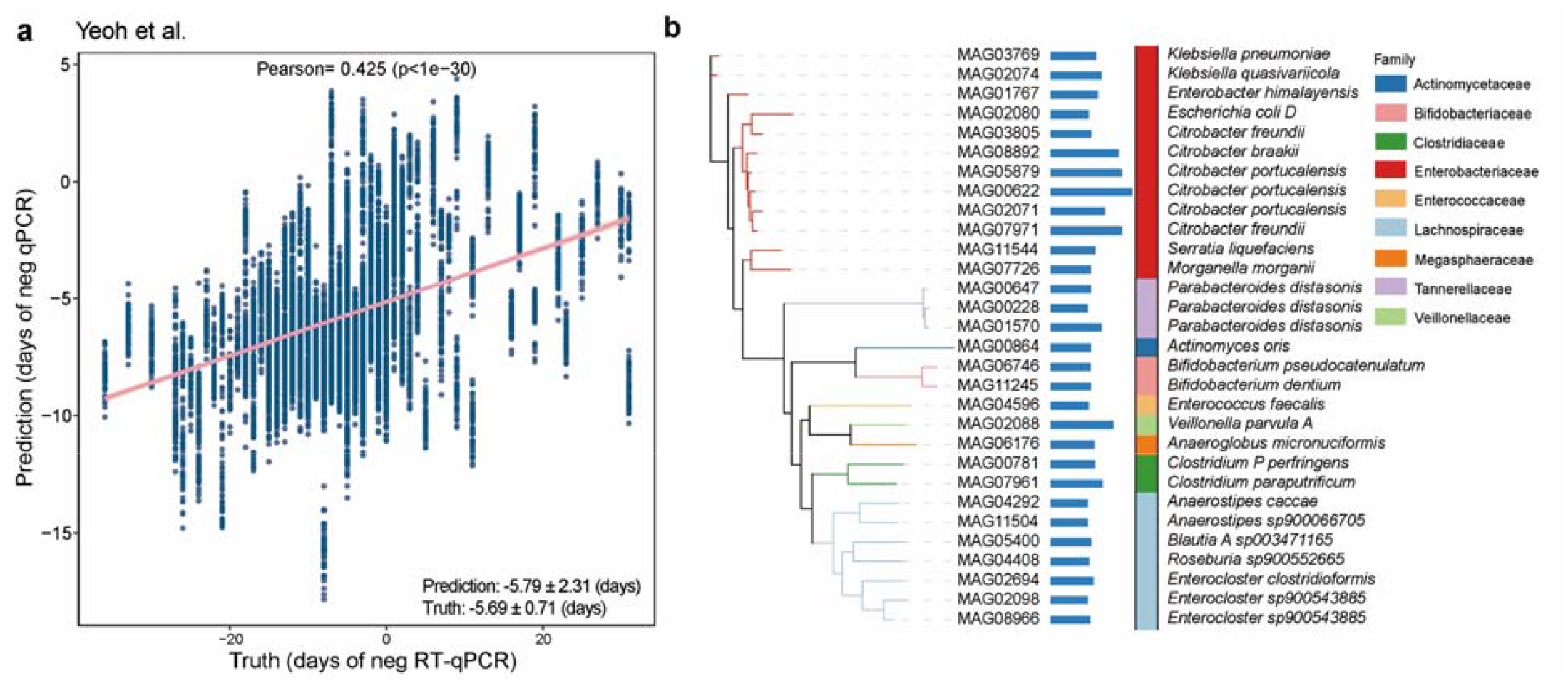
The nrMAG-based machine learning model predict the progression of COVID-19. **a**, Pearson correlation coefficient between the true and predicted date of negative RT-qPCR result on the random forest regression models. **b**, The top 30 important nrMAGs related to performance of prediction. The importance of each feature in the regression was quantified by percent increase in mean square error (%IncMSE). The length of horizontal bar represents the mean %IncMSE value. The colors of vertical bar represent the taxonomy information of nrMAGs at the family level. The phylogenetic tree of these nrMAGs was constructed using PhyloPhlAn.

### Identification of putative permissive and protective nrMAGs for COVID-19 severity

To further characterize the relation between the human gut microbiome and COVID-19, we applied the generalized microbe-phenotype triangulation (GMPT) method to move beyond the standard association analysis^44^ (**Fig.5a**). We first categorized participants into five different disease severity groups (i.e., Non-COVID-19 healthy controls, mild, moderate, severe, and critical) using data from Yeoh et al.^18^ The differentially abundant nrMAGs were then calculated using ANCOM^45^ in the ten pairwise comparisons. Using this approach, all pairwise differential abundance analyses yielded a total of 644 differentially abundant nrMAGs present in at least two pair-wise comparisons. To understand the potential relationship between those candidate nrMAGs with COVID-19, we then calculated the Spearman correlation coefficients between the average relative abundances and COVID-19 severity score (e.g., Non-COVID-19 healthy controls: 0; mild: 1; moderate: 2; severe: 3; and critical: 4) in different phenotype groups. Those differentially abundant nrMAGs with positive (or negative) Spearman correlation coefficients are potential permissive (protective) nrMAGs of COVID-19. Based on the frequency (≥ 6) of all pairwise comparisons (n=10), we summarized the results from GMPT in **Fig.5b** and **Table S1**. This analysis identified a total of 74 nrMAGs that were associated with SARS-CoV-2 infection, including 8 permissive (Spearman correlation > 0) nrMAGs, 63 protective (Spearman correlation < 0) nrMAGs, and 3 neutral nrMAGs (Spearman correlation = 0).

**Fig. 5.**
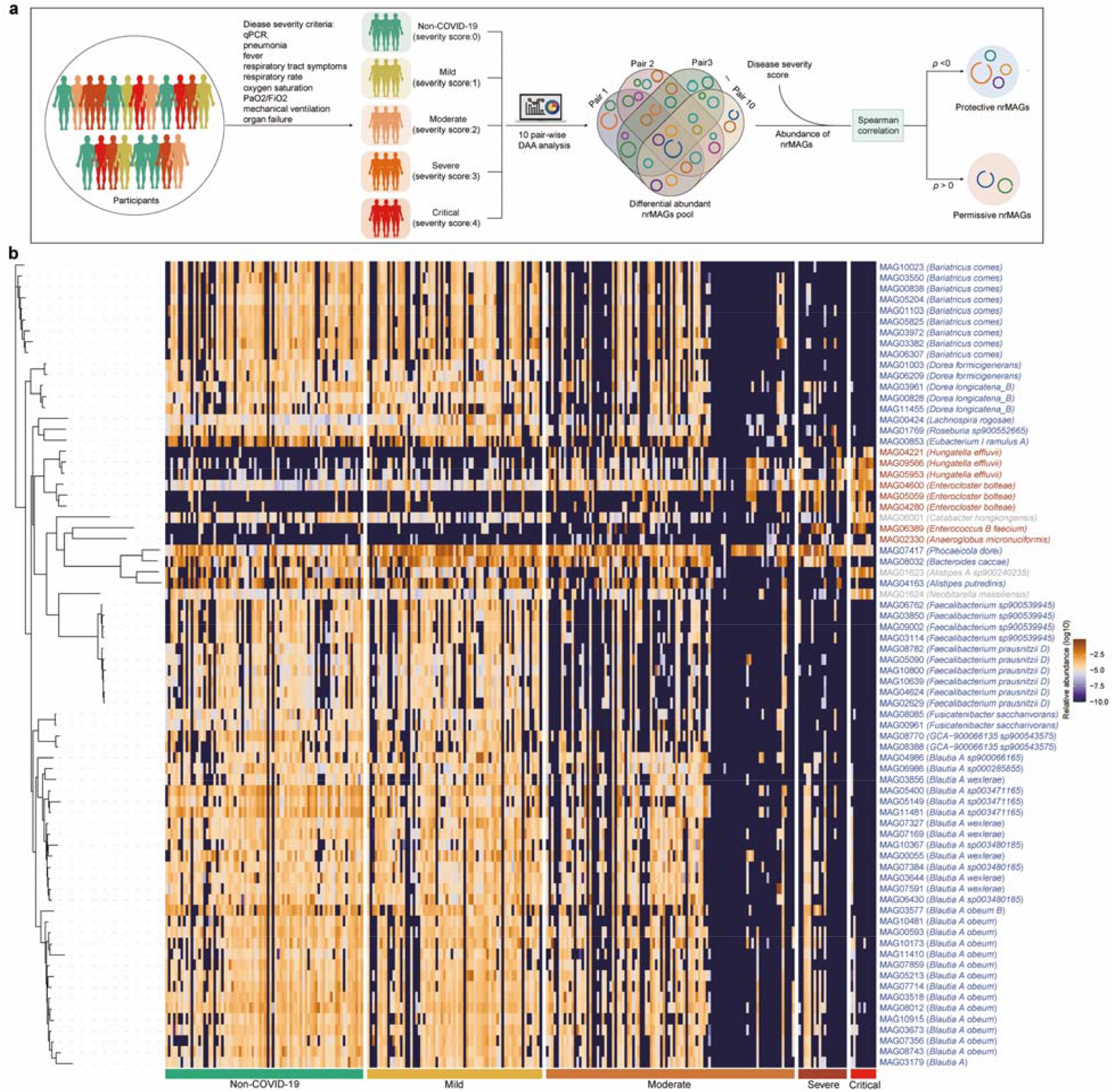
The permissive and protective nrMAGs of COVID-19 identified by GMPT pipeline. **a, The workflow of GMPT pipeline**. The microbiome samples from the study of Yeoh et al. grouped to 5 groups (i.e., healthy control, mild, moderate, severe, and critical) based on disease severity. Differential abundance analysis was carried out on each possible pairwise comparison (10 pair wise comparisons among 5 groups). The differentially abundant nrMAGs pool were originated from these pairwise analyses. The differentially abundant nrMAGs were ranked based on their frequency appearing in all the pairwise comparisons and differentiability (descending order). We further used the average relative abundance of nrMAGs and the disease severity to calculate the Spearman correlation coefficient. Here positive (negative) Spearman correlation coefficient (ρ) represent the permissive (protective) nrMAGs of COVID-19 severity. And the spearman correlation with 0 means the nrMAGs may be neutral to the severity of SARS-CoV-2 infection. **b**, The heat map showed the abundance distribution of permissive, neutral, and protective nrMAGs across different disease severity groups identified using GMPT. These nrMAGs were taxonomically annotated using GTDB-Tk based on the Genome Taxonomy Database. The colors of the taxonomical label represent permissive, protective, or neutral nrMAGs. The phylogenetic tree of these nrMAGs was constructed using PhyloPhlAn.

We identified multiple species with highly similar genomes from those permissive nrMAGs (**Fig.5b**), including *Enterocloster bolteae* (3 nrMAGs), *Anaeroglobus micronuciformis, Hungatella effluvii* (3 nrMAGs), and *Enterococcus B faecium*. Consistent with previous reports that the gut microbiome of COVID-19 patients showed significant higher abundance of *Enterococcus faecium* compared to health controls^46^. Among the 63 protective nrMAGs, the dominant species were *Blautia A obeum* (13 nrMAGs), *Bariatricus comes* (9 nrMAGs), *Faecalibacterium prausnitzii D* (6 nrMAGs), *Blautia A wexlerae* (6 nrMAGs), *Faecalibacterium sp900539945* (4 nrMAGs), *Dorea longicatena B* (3 nrMAGs), *Blautia A sp003480185* (3 nrMAGs), *Blautia A sp003471165* (3 nrMAGs), *Dorea formicigenerans* (2 nrMAGs), *Fusicatenibacter saccharivorans* (2 nrMAGs), and *GCA-900066135 sp900543575* (2 nrMAGs). Importantly, we found that some of these species were previously reported (including in the original study Yeoh et al.) to be decreased in patients with COVID-19 such as *Blautia obeum*^11,18^, *Faecalibacterium prausnitzii*^17,18,39,47^, and *Dorea formicigenerans*^17,18^. Interestingly, those protective nrMAGs also showed a similar abundance distribution between patients with COVID-19 and Non-COVID-19 controls in the study of Zuo et al^17^ (**Fig.S9**). This finding provides strain level evidence that gut microbial taxa may interact with SARS-COV-2 infection and play a potential role in disease onset and progression in COVID-19.

### Genome annotation reveals functional differentiation between permissive and protective nrMAGs of COVID-19

Information on biological products encoded in the human microbiome will facilitate the functional characterization of disease-associated microbiota. This prompted us to investigate whether the functional capacity of permissive and protective nrMAGs differ. To achieve that goal, we first annotated the genomes of permissive and protective nrMAGs using Prokka^48^. Then we processed the translated coding sequences using MicrobeAnnotator for the functional annotation and calculated the KEGG module completeness (see Methods)^49^. Here, KEGG modules are functional gene units, which are linked to higher metabolic capabilities, structural complexes, and phenotypic characteristics^49^. A total of 231 and 254 KEGG modules were covered by at least one genome from permissive and protective nrMAGs, respectively. Principal component analysis revealed quite different metabolic potentials between permissive and protective nrMAGs (**Fig.6a**, PERMANOVA: *P* value =0.0001). The main KEGG modules (with at least 50% module completeness) of each nrMAG are summarized in **Fig.S10**. Notably, we identified a set of KEGG modules that differed significantly in their module completeness between permissive and protective nrMAGs (**Fig.6b**). For example, permissive nrMAGs showed significantly higher completeness level at the pentose phosphate pathway (e.g., M0004 and M0006) compared to protective nrMAGs. Importantly, a previous study reported a significant increase in the levels of some intermediates of the glycolytic and pentose phosphate pathways in sera of COVID-19 positive patients^50^. Moreover, SARS-CoV-2 infection was found to be associated with changes in the regulation of the pentose phosphate pathway in both in vivo (Caco-2 cells)^51^ and in vitro (ferret model)^52^ studies. These findings provide evidence of potential metabolic pathways linking the gut microbes to the clinical manifestations of COVID-19.

**Fig. 6.**
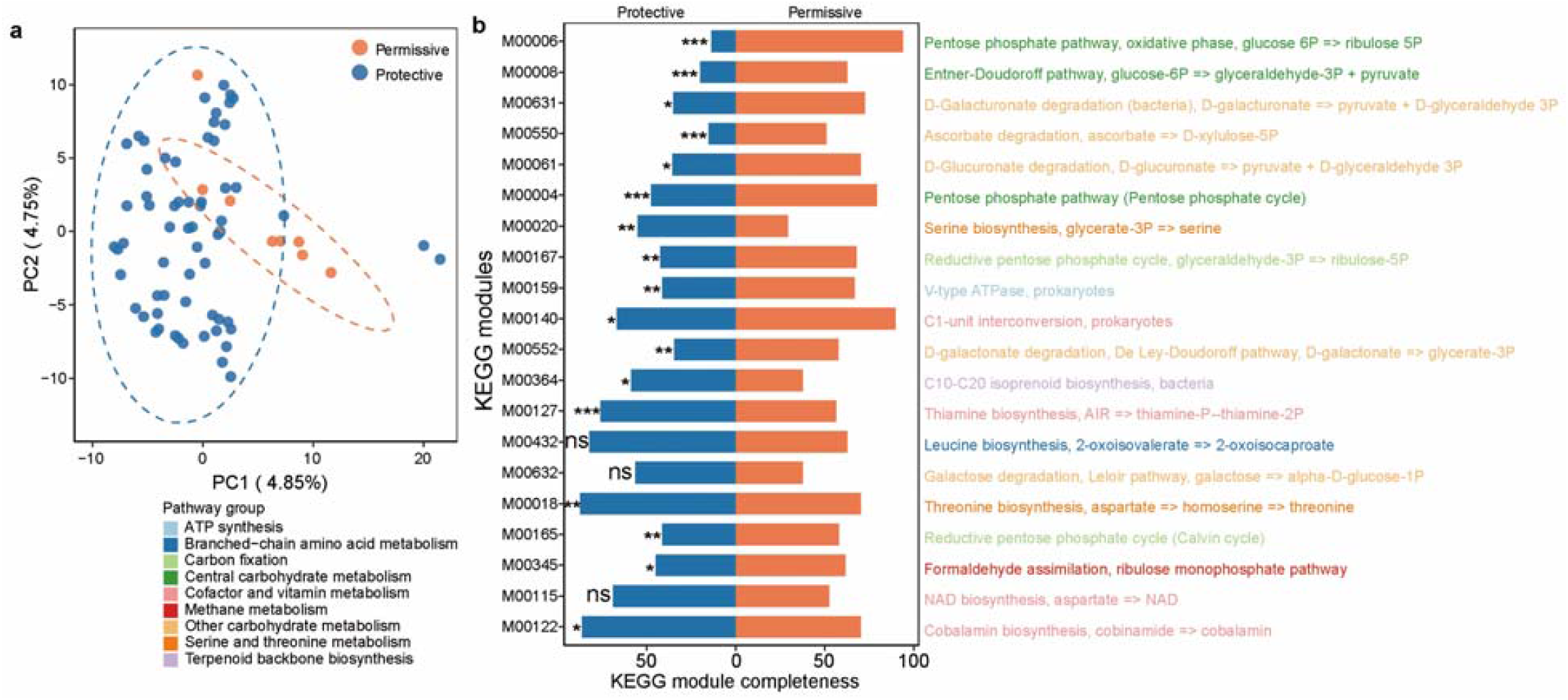
Genome annotation of permissive and protective nrMAGs of COVID-19. **a**, Principal Component Analysis (PCA) plot of KEGG module completeness from all genomes of permissive and protective nrMAGs. **b**, The top 20 differential KEGG modules between permissive and protective nrMAGs ranked based on the mean difference. *P* values were calculated by two-sided Wilcoxon–Mann–Whitney test (ns: nonsignificant; *: *P*<0.05; **: *P*<0.01, ***: *P*<0.001).

## Discussion

Here, we leveraged a total of 514 publicly available WMS sequenced samples from 6 SARS-COV-2 data sets and generated for the first time a high-quality COVID-19 related genome catalog of the human microbiome. We recovered a large genome catalog representing 11,584 MAGs and 5,403 nrMAGs of the human microbiome. Through the construction of this microbial genome catalog, we were able to provide the strain-level perspective to understanding the human microbiome and COVID-19.

By interrogating the WMS sequencing data with different technical settings, we gained a more comprehensive view of the microbial community associated with COVID-19. Due to the inherent differences (e.g., age, diet, and genetic background) across the different data sets, our goal was not targeted comparisons across data sets. Importantly, our study mainly focused on two data sets (Zuo et al.^17^ and Yeoh et al^18^.) with well-defined case and control subjects. Although two nasopharyngeal microbiome data sets contained both patients with COVID-19 and Non-COVID-19 controls, the statistical power was limited by the small sample size (Liu et al.^35^) and a large portion of sequencing reads from another data set (PRJNA743981^36^) were from the human host. Our analysis did not exclude these nasopharyngeal microbiome samples as they did contribute unique high-quality MAGs to our nrMAGs collection. Furthermore, two gut microbiome data sets (Britton et al.^53^ and Cao et al.^54^) without Non-COVID-19 controls served as key resources of our MAGs collections.

Previous efforts have linked human microbiome diversity and COVID-19^11,14,33^. Coherently, the human gut microbiome of patients with COVID-19 in our study exhibited decreased alpha diversity at the nrMAGs level compared to the Non-COVID-19 healthy controls. Although we did not found significant differences between patients with COVID-19 and Non-COVID-19 controls in two nasopharyngeal microbiome data sets, this may be due to the small sample size and the fact that Non-COVID-19 controls are not health controls^35^. Notably, our analysis identified that patients with COVID-19 after recovery (negative for SARS-CoV-2 via RT-qPCR) differed more from Non-COVID-19 controls compared to patients with COVID-19 before recovery. This finding supports the possibility that the gut microbiome of patients with COVID-19 may not return to a relatively healthy status right after their recovery from COVID-19^55^. Given the fact that many patients recovering from SARS-CoV-2 infection have experienced prolonged COVID-19 symptoms^56^, we hypothesize that long-lasting disease symptoms may be associated with changes in the gut microbiome but this needs to be explored further. Our analysis also supports the main findings from the original study of Yeoh et al^18^. That the gut microbiota composition reflects disease severity. Specifically, we revealed this connection through the Bray-Curtis dissimilarity comparison between severity group and healthy controls, and changes in microbial community membership longitudinally overtime at the nrMAGs level. Interestingly, the connection between disease severity of COVID-19 and the gut microbiome has also been reported at both the species- and metabolic capacity-level by a recent study^39^.

Using machine learning models, we demonstrated that the gut microbiome signatures at the nrMAG-level can accurately detected COVID-19 from healthy controls. The high diagnostic accuracy of our microbiome-derived signature suggests that key microbial strains within the signature might play important roles in the pathogenesis of COVID-19. For example, some of nrMAGs at higher taxonomic levels (e.g., genus and species) have been reported to be correlated with COVID-19, such as *Blautia*^11,15,18^, *Faecalibacterium prausnitzii*^17,18,47^, and *Adlercreutzia equolifaciens*^18,21^. Notably, this study sheds important light on the ability of nrMAGs to predict the date of negative RT-qPCR result of patients with COVID-19. This analysis linked several microbial species from our important nrMAGs to the progression of COVID-19 such as *Citrobacter freundii, Veillonella parvula*, and *Parabacteroides distasonis*. Indeed, these species have been previous reported involved in COVID-19. For example, *Citrobacter freundii* was found to be significantly enriched in COVID-19 patients with fever^57^. *Veillonella parvula*^18,40,54^ and *Parabacteroides distasonis*^18,58^ were also shown to be a shared signature of COVID-19 in multiple studies. Importantly, we observed some opportunistic pathogens were associated with the progression of COVID-19, including nrMAGs from *Klebsiella quasivariicola*^41^, *Klebsiella pneumoniae*^42^, and *Escherichia coli*^43^. Related to our findings, multiple studies revealed high prevalence of bacterial pathogens in patients with COVID-19^17,59-62^, further supporting the possibility that secondary infections by opportunistic pathogens may affect the progression of COVID-19.

In particular, host gut microbiota provides colonization resistance against pathogens, for example, a previous study reported that mice treated with neomycin antibiotics were more susceptible than control mice to influenza viruses^63^. And it turned out that neomycin-sensitive bacteria naturally present in the mice’s bodies provided a trigger that led to the production of T cells and antibodies that could fight an influenza infection in the lungs. Notably, our study identified a set of COVID-19 related nrMAGs and their determinants (i.e., permissive, and protective) potentially involved in disease pathogenesis. It is important to note that these protective bacteria have also been reported to be related to SARS-CoV-2 infection. For example, the host immunity symbionts beneficial bacterial species *B. obeum* was identified to be depleted in patients with COVID-19 in multiple studies^11,17^. *F. prausnitzii*, an anti-inflammatory and butyric acid-producing commensal bacterium, was found to be underrepresented in patients with COVID-19^17,18,64^. Moreover, our analysis enables the direct microbial genome comparison between permissive and protective nrMAGs. Consistent with previous reports of a relationship of the pentose phosphate pathway and SARS-CoV-2 infection^50-52^, our findings support that overrepresentation of permissive nrMAGs and underrepresentation of protective nrMAGs may upregulate the pentose phosphate pathway as their genome are shown to be highly intact in those relevant modules. In contrast, the pentose phosphate pathway related KEGG modules showed significant lower module completeness in genomes of protective nrMAGs. Together, these results provide valuable insights into the mechanisms underlying host-microbiome interaction and COVID-19 severity and progression, and most importantly, future work may then be able to capture molecular mechanisms of these specific strains more easily in the severity of SARS-CoV-2 infection.

The current study has several limitations. First, although we included a large number of shotgun metagenomic sequencing samples from the COVID-19 related human microbiome study (publicly available as of August 2021), most of the microbiome samples came from China. This limitation could be addressed by following this work with collection of more human microbiome samples from different populations and body sites to construct a more comprehensive genome catalog to reveal the full landscape of the human microbiome in COVID-19. Second, even though we adjusted for some potential confounders in our statistical models, we were unable to assess some covariates such as: medication, diet, and psychological stress that are not publicly available. Third, consistent with multiple MAG-related WMS studies^25,65,66^, we only recovered MAGs from bacteria and archaea. Given the fact that *de novo* discovery of non-bacterial genomes is quite challenging^67^, future study targets for other domains, including fungi and viruses, will give a more comprehensive view in the context of host-specific microbiotas and COVID-19. Although the majority of MAGs we reconstructed in this study have high quality, future investigations aiming for recovering the complete genome of microbes will further enhance our understanding of the interaction between human microbiome and SARS-CoV-2 infection^68,69^. Finally, additional experiments are needed to assess the casual role of candidate permissive and protective nrMAGs in COVID-19 progression. Nevertheless, given the large uncultured diversity still remaining in the human gut microbiome and deficiency of both annotated genes and reference genomes, having a high-quality genome catalog substantially enhances the resolution and accuracy of metagenome-based COVID-19 studies. Therefore, the presented genome catalog represents a key step toward mechanistic understanding the role of the human gut microbiome in SARS-CoV-2 infection.

In summary, we present here the first construction of the genome catalog using assembly and reference free binning of metagenome in patients with COVID-19 and Non-COVID-19 controls. Our findings support the close connection between SARS-CoV-2 infection and the human gut microbiome. These insights into metagenomic strain-level aspects of relation in human microbiome and COVID-19 and genome context will form the basis of future studies.

## Acknowledgements

We would like to thank Dr. Yun Kit Yeoh and Dr. Siew C Ng for sharing the phenotypic data with us. We thank Xu-Wen Wang, Zheng Sun, Tong Wang, Darius Schaub, Yunyan Zhou, and Xiaochang Huang for valuable discussions. Yang-Yu Liu acknowledges the funding support from the National Institutes of Health (R01AI141529, R01HD093761, RF1AG067744, UH3OD023268, U19AI095219, and U01HL089856).

## Author contributions

S.K. and Y.-Y.L conceived and designed the project. S.K. performed all the data analysis. S.K. and Y.-Y.L interpreted the results and prepared the manuscript. S.T.W reviewed and edited the manuscript. All authors approved the manuscript.

## Methods and materials

### Data collection

We identified COVID-19 metagenomic sequencing studies from keyword searches in PubMed and online repositories (i.e., NCBI, ENA, and GSA) and by following references in meta-analyses and related microbiome studies. We included samples with publicly available raw shotgun metagenomic sequencing data (paired fastq files) and metadata indicating patients with COVID-19 or Non-COVID-19 control status. All the sequencing data were downloaded from online repositories or links provided in the original publications, but some metadata were acquired after personal communication with the authors. We did not include any studies which required additional ethics committee approvals or authorizations for access. A total of 514 microbiome samples from six public microbiome data sets were analyzed in this study (Table 1)

### Metagenome assembly and binning

Genome reconstruction of human microbiome with metagenomic sequencing data was performed with the function modules of metaWRAP (v1.3.2)^70^, which is a pipeline that includes numerous modules for constructing metagenomic bins. First, the metaWRAP-Read_qc module was applied to trims the raw sequence reads and removes human contamination for each of the sequenced samples. Then the clean reads from the sequencing samples were assembled with the metaWRAP-Assembly module using metaSPAdes (v3.13.0)^71^. Thereafter, MaxBin2 (v2.2.6)^72^, metaBAT2 (v2.12.1)^73^, and CONCOCT (v1.0.0)^74^ were used to bin the assemblies. The default of the minimum length of contigs used for constructing bins with MaxBin2 and CONCOCT were 1,000 bp, and metaBAT2 was defaulted to 1,500 bp^70^. Refinement of MAGs was performed by the bin_refinement module of metaWRAP^70^, and CheckM (v1.0.12)^75^ was used to estimate the completeness and contamination of the bins, and the minimum completion and maximum contamination were 50% and 10%, respectively.

### Species-level clustering and dereplication and of MAGs

All 11,584 MAGs were clustered into species-level genome bins (SGBs) at the threshold of 95% ANI using the ‘cluster’ program in dRep (v3.0.0)^76^. All MAGs were taxonomically annotated using GTDB-Tk (v.1.4.1)^77^ based on the Genome Taxonomy Database (http://gtdb.ecogenomic.org/)^29^, which produced standardized taxonomic labels that were used for the analysis in this study. SGBs containing at least one reference genome (or MAG) in the Genome Taxonomy Database were considered as known SGBs. And SGBs without reference genomes were considered as unknown SGBs (uSGBs)^67^. dRep (v3.0.0)^76^ was then used for dereplication of all 11,584 MAGs (≥50% genome completeness and ≤5% contamination) by two-steps. First, MAGs were divided into primary clusters using Mash^78^ at a 90% Mash ANI. Then, each primary cluster was used to form secondary clusters at the threshold of 99% ANI with at least 30% overlap between genomes. According to the criteria of quality evaluation by CheckM (v1.0.12)^75^, 5403 nrMAGs were divided into medium-quality MAGs (50% ≤ completeness < 90% and ≤5% contamination) and high-quality MAGs (≥90% completeness and ≤5% contamination).

### Abundances estimation and phylogenetic analysis of nrMAGs

The metaWRAP-Quant_bins module integrated with Salmon^79^ (v0.13.1) was used to estimate the abundance of each nrMAGs in each of the metagenomic samples. The phylogenetic tree of the nrMAGs was built using PhyloPhlAn (v3.0.58)^80^. The tree was visualized using iTOL (https://itol.embl.de/)^81^.

### Genome annotation of MAGs

The genome annotation of MAGs was first performed with Prokka (v1.13)^48^ using the annotate_bins module of metaWRAP^70^. The annotated genomes were then processed with MicrobeAnnotator (v2.0.5)^49^ for the functional annotation and to calculate KEGG module completeness. All proteins are searched against the curated KEGG Ortholog (KO) database using Kofamscan^82^; best matches are selected according to Kofamscan’s adaptive score threshold. Proteins without KO identifiers (or matches) are extracted and searched against other databases (e.g., Swissprot, curated RefSeq database or non-curated trEMBL database)^49^. The KO identifiers associated with all proteins in each genome (or set of proteins) are extracted, and KEGG module completeness is calculated based on the total steps in a module, the proteins (KOs) required for each step, and the KOs present in each genome. Finally, the results were compiled in a single matrix-like module completeness table for all genomes.

### Statistical analysis

Microbial alpha and beta diversity measures were calculated at the nrMAGs level using vegan package (v2.5.7) in R. Principal coordinates analysis (PcoA) plots were generated with Bray– Curtis dissimilarity. Differences in microbiome compositions across different groups were tested by the permutational multivariate analysis of variance (PERMANOVA) using the “adonis” function in R’s vegan package. All PERMANOVA tests were performed with 9999 permutations based on Bray–Curtis dissimilarity. Differences between groups were analyzed using a Wilcoxon–Mann–Whitney test. For differential abundance analysis in GMPT (Generalized Microbe Phenotype Triangulation) pipeline^44^, we used ANCOM (analysis of composition of microbiomes)^45^, with a Benjamini–Hochberg correction at 5% level of significance, and adjusted each patient’s identifier as a random effect. Only the nrMAGs that were presented in at least 5% of samples were included. The phylogenetic tree of the permissive, neutral, and protective nrMAGs was built using PhyloPhlAn (v3.0.58)^80^ and then visualized using iTOL (https://itol.embl.de/)^81^.

To develop a model capable of distinguishing patients with COVID-19 versus Non-COVID-19 controls, we implemented Random Forest using R’ random Forest package. A custom machine learning process was conducted using features of nrMAGs with 5-fold cross validation. The data was split into a training set and a test set, with 80% of the data forming the training data and the remaining 20% forming the test set. And then we randomly split the data 50 times. The performance of the classification model was evaluated using AUROC (area under the receiver operating characteristic curve) and AUPRC (area under the precision-recall curve) on the test set. The importance of each feature was quantified by the Mean Decrease in Accuracy (MDA) of the classifier due to the exclusion (or permutation) of this feature. The more the accuracy of the classifier decreases due to the exclusion (or permutation) of a single feature, the more important that feature is deemed for classification of the data. We then built Random Forest regression model with 5-fold cross-validation to predict the exact date of microbiome sample collected before or after negative RT-qPCR result. We randomly split the data 50 times. The importance of each feature in the regression was quantified by percent increase in mean square error. Pearson correlation coefficient between the true and predicted date of negative RT-qPCR result was used to evaluate the performance. All statistical analysis was performed with R (version 3.6.3).

### Data availability

All data used in this article come from publicly available sources. The accession numbers of those metagenomic studies are PRJNA624223, PRJNA656660, PRJNA660883, PRJNA743981, PRJCA003532, and PRJNA650244.

